# Improving SCVI for low-count cells through self-supervised augmentation

**DOI:** 10.64898/2026.02.11.705441

**Authors:** Valentine Svensson

## Abstract

When analyzing single-cell RNA sequencing data with SCVI, low-UMI cells typically need to be filtered, because their learned representations lack meaningful biological signal. We show that this is caused by a specific mechanism: as UMI depth decreases, the SCVI encoder maps cells towards a learned bias point, collapsing their representations regardless of cell identity. This phenomenon is distinct from classical posterior collapse driven by KL regularization. By augmenting training with binomial thinning and adding a cross-correlation loss between original and thinned cell embeddings, the encoder learns representations that preserve cell type identity, experimental condition differences, and sample-level variation at lower UMI depths, without sacrificing reconstruction quality. These modifications extend the range of usable cells, enabling analysis of cells that would typically be discarded due to low molecule counts.

---

Single-cell RNA sequencing (scRNA-seq) is a technology which allows researchers to investigate which cells are expressing which genes, and under which circumstances. It has enabled the discovery and cataloging of distinct cell types across all areas of biology (Aldridge & Teichmann, 2020).

Variation in the total molecule counts (‘library size’) between cells have been identified as a strong source of nuisance variation in scRNA-seq data (Hicks et al., 2018; Lause et al., 2021; Townes et al., 2019).

Making use of a count-based variational autoencoder, the SCVI model is designed to integrate out variation in library size, both in low-dimensional representations and in normalized expression levels on a relative abundance scale (Gayoso et al., 2022; Lopez et al., 2018).

While the SCVI model alleviates a large amount of spurious correlations due to variations in library size, cells with extremely low depth often separate from cells with high total counts in their learned representations^1^.

Cells with low total UMI content (unique molecular identifiers; deduplicated molecule counts) provide less information about the relative mRNA abundance from different genes. To focus analysis on more informative cells, it is recommended to filter out cells with particularly low total UMI content (Heumos et al., 2023; Luecken & Theis, 2019).

The number of cells that need to be filtered out is specific per dataset, and can vary depending on how challenging it is to isolate the cells, the efficiency of molecular reagents, and sequencing depth, among other factors (Ding et al., 2020; Svensson et al., 2019).

Enabling successful analysis of cells with lower total UMI counts would provide more utility from precious tissue samples with isolation challenges. It would increase the amount of useful data using more cost-efficient reagents and lower sequencing budget.

To address this, we used count subsampling through binomial thinning on six typical scRNA-seq datasets to identify the mechanism by which the SCVI model starts separating cells with low UMI content. This insight was used to modify the loss function and training procedure of the SCVI model for improved performance on cells with lower counts.

As total cell UMI counts decrease, the cells converge towards a learned bias point implicit in the SCVI encoder. Modifying the training procedure to perform binomial thinning of the cells and adding a cross-correlation loss with original cells decreases the rate of convergence to the bias point (Zbontar et al., 2021). As a consequence, the encoder learns to retain biologically meaningful cell type identities and experimental effects for cells with lower count depths.

With these modifications, a larger fraction of cells can be used for analysis, which is particularly important for precious samples such as archival tissues where cell isolation can be challenging. It also increases the amount of useful data for training foundation models. Similar training and loss strategies could be applied to other single cell model families and tasks.

## As library size depth decreases, cells converge towards a learned bias point

When visualizing or clustering single cells modeled by SCVI, it is typical to find that low-UMI cells are distant from high-UMI cells. Cells with very low total UMI counts group together, with no clear cell type identity.

To improve our ability to model cells at lower counts we need to understand the mechanism behind this phenomenon. We investigated this by artificially reducing the total UMI counts of high-UMI cells through binomial thinning, then passing the cells through a trained SCVI encoder. Results were compared across six different representative datasets from different biological systems.

During training, the inference model (encoder) of the SCVI model learns to encode a *bias point*, representing a cell with zero observed molecules from each gene.

As UMI depth of cells decreases, they approach this bias point, converging on it at extremely low UMI depths.

### Low-UMI bias point convergence is unrelated to KL-driven posterior collapse

Kullback-Leibler (KL) regularization is designed to pull representations of low-information observations towards the prior, centered at the origin (P Kingma & Welling, 2019). A common failure mode of variational autoencoders is *posterior collapse* where latent representations are uninformed by the encoded data, and instead are sampled from the prior, often attributed to the KL loss term (James et al., 2019). This is particularly relevant for the negative-binomial log-likelihood loss in SCVI models, since the KL term will have higher relative weight for low-UMI cells than for high-UMI cells.

However, the learned bias point can be as far away from the origin as representations of high-UMI cells (Figure 1).

**Figure 1:**
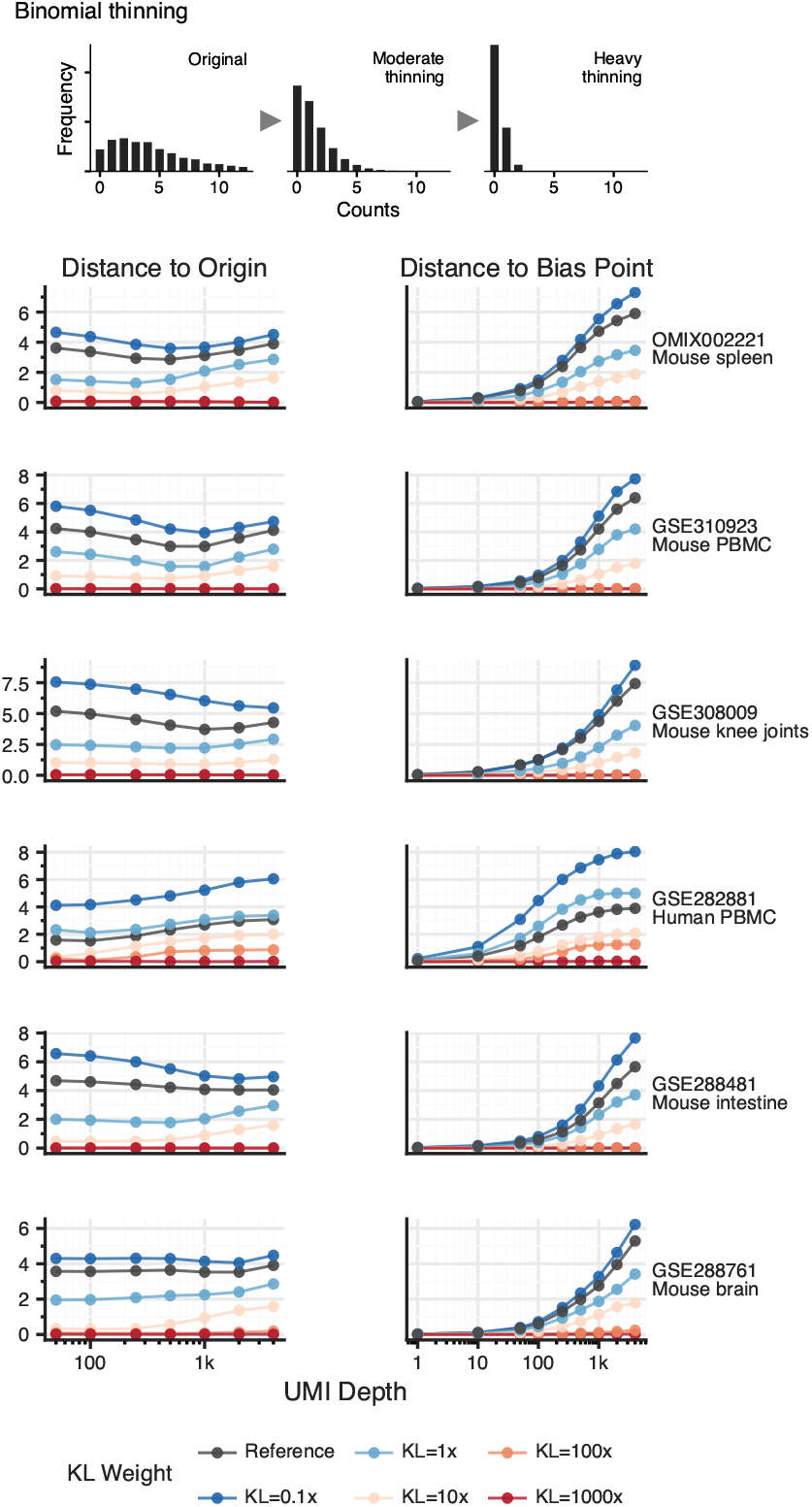
Behavior of cell representations from SCVI encoders after binomial thinning of high-UMI cells.

Massively increasing the contribution of the KL term to the loss recovers the traditional posterior collapse phenomenon (Figure 1, Figure S1, Table S1).

Low-UMI cells converging to a learned bias point is a distinct phenomenon from posterior collapse.

To make better use of low-UMI data, we need to identify modeling strategies which discourages the encoder from learning to move cells towards the bias point as total UMIs decrease.

## Addition of a cross-correlation loss leads to better low-UMI cell representations

The standard SCVI encoder increasingly mixes cells together as total UMI decreases. To reduce sensitivity to UMI depth in the encoder, we introduced self-supervision through binomial thinning augmentation during training, along with a cross-correlation loss term.

Using binomial thinning of high-UMI cells, we show that this training strategy and cross-correlation loss produces better representations of low-UMI cells.

The added loss and training strategy decreases the rate at which cells converge to the bias point. As a consequence, the encoder maintains cluster membership at lower depths, and better preserves the ability to distinguish cells from different experimental conditions (Figure 2). Adding this cross-correlation objective during training has minimal impact on reconstruction quality (Table S2).

**Figure 2:**
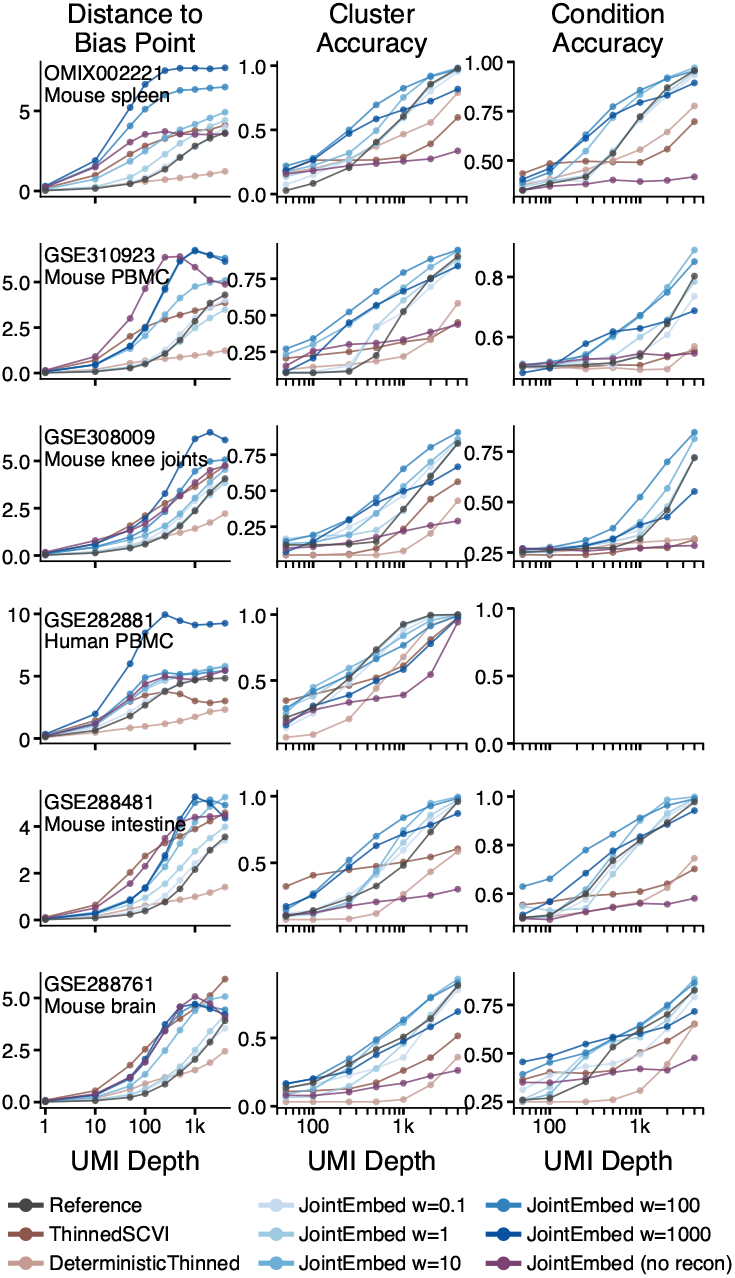
Behavior of cell representations, and performance on biologically motivated tasks on cells representations, after binomially thinning high-UMI cells and passing them through model encoders.

These training modifications of SCVI models lead to encoders that are more reliable at lower UMI depths. Using models trained with the cross-correlation objective will allow higher resolution at low depths, and the ability to make use of cells that are typically filtered out from analysis.

### Data augmentation alone is insufficient

The addition of a loss term adds complexity, in particular regarding balancing the terms. As an alternative, we modified the training procedure with binomial thinning before encoding, maintaining the standard loss terms.

Training with data augmentation alone led to worse performance in biological tasks, even at high UMI depths (Figure 2).

Cross-correlation loss is needed for the encoder to learn useful representations.

### Gene expression reconstruction loss is necessary

To encode cell representations reflecting biological states, reconstruction of gene expression might not be needed, substantially simplifying the model. We trained models with pure self-supervision using the cross-correlation objective, without reconstruction.

Self-supervision alone results in encoders that avoid converging on the bias point at lower UMI depths, but they are unable to reflect cluster membership or separation between experimental conditions (Figure 2, Table 1).

**Table 1:**
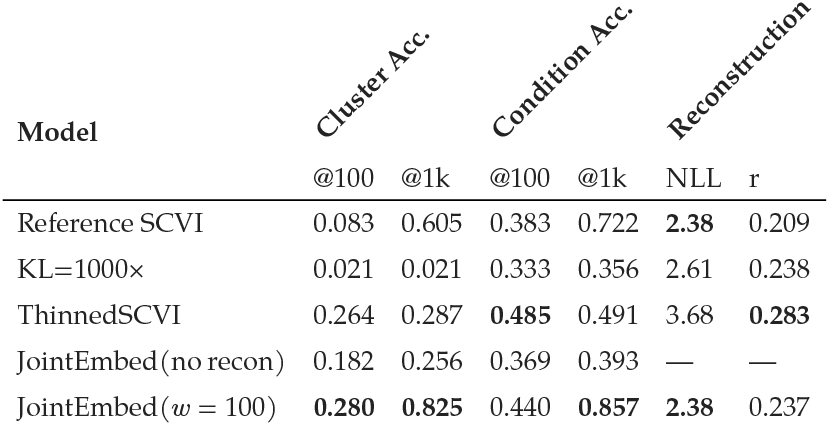
Representative ablation results on OMIX002221 (Mouse spleen). Cluster accuracy and condition accuracy at binomially thinned 100 and 1000 UMI depth, and representative reconstruction metrics (negative log likelihood and Pearson correlation on non-zero entries).

Without the model tying cell representations to gene expression output, biological signals in the data are lost.

## Conclusions

When analyzing scRNA-seq data, strong filters for a minimal number of total UMIs are needed when using SCVI models. In standard SCVI models, low-UMI cells converge to a learned bias point, a phenomenon that is unrelated to the classical posterior collapse in variational autoencoders.

By modifying the training procedure and adding a cross-correlation loss when training SCVI models, the encoders learn to represent meaningful biological states for cells with lower UMI depth.

Without sacrificing reconstruction quality, these modifications enable analysis including many low-UMI cells that are typically filtered out before analysis.

This is particularly important for archival tissue samples, where extracting cells with high UMI counts is challenging. It also enables analysis of data generated using less efficient reagents and lower sequencing budgets.

Reconstruction (for normalization and differential expression analysis) and representation (for cell state analysis, clustering analysis, featurization for classification) are common tasks SCVI models are used for. Another common task is integration, where variation due to known factors (such as technical batches or patients) are integrated out from learned representations. Integration performance is not considered here, and it is not clear how cross correlation loss affects integration performance.

## Methods

### The SCVI model for scRNA-seq data

The generative model of SCVI can be defined as

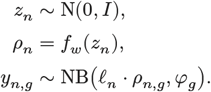

This generative model is paired with an inference model (encoder):

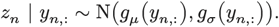

The model is trained using the combined loss

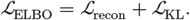

In all experiments, we set *D* = 32 based on recent hyper parameter study (Kassab et al., 2026), and trained the model for 25 epochs with a batch size of 512 and learning rate of 0.004^2^. We also disable KL warmup, allowing full KL regularization throughout training.

### Scaling the KL term to the number of genes

The reconstruction loss term in the SCVI model is the negative log likelihood

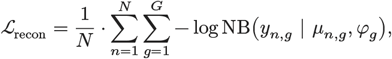

where *µ*_*n,g*_ = *ℓ*_*n*_ *⋅ ρ*_*n,g*_ are model predictions. This loss measures how compatible observed UMI counts from the genes in the cells are to a statistical generative model of the count data. It combines the negative log likelihoods of the genes to provide an average per-cell measure.

On the other hand, the KL loss term is

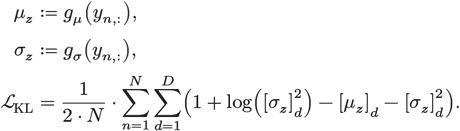

Comparing the two terms, ℒ_recon_ sums over the genes, while ℒ_KL_ sums over the latent dimensions. This means models fitted with different datasets that have different numbers of genes will exhibit different levels of influence from the KL regularization.

To make model behaviors comparable between individual datasets, we scale the KL term according to

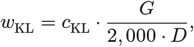

where, in most cases, *c*_KL_ = 1. The constant 2,000 is motivated by the original KL divergence strength typically being applied to ∼2,000 representative genes used for analysis (Lopez et al., 2018).

### Reconstruction-based self-supervised learning

The SCVI model is a probabilistic generative model of UMI counts where the variational posterior of latent variables can be used as cell representations for downstream tasks.

Aside from the probabilistic framing, we can view the model as a reconstruction-based self-supervised representation learning model. The SCVI model performs data augmentation in the latent space during training through *z*_*n*_ = *g*_*µ*_(*y*_*n*,:_) + *g*_*σ*_(*y*_*n*,:_) *⋅ ε*, where *ε ∼* N(0, *I*).

### Data augmentation for scRNA-seq

In a given cell from scRNA-seq data we observe molecule counts from each gene which are proportional to the abundance of mRNA from the genes in the cell. There are often biological reasons for why different cells have different total mRNA abundances, but such differences will *also* be reflected in differences of relative abundances between mRNA levels from different genes. To investigate biological variation, differences in total mRNA abundances (without differences in relative abundances) are often of little interest. In most situations, differences in total abundance is a technical, statistical, or economic factor.

Observed counts in scRNA-seq data follow count statistics, adhering to the sampling properties that arise as exposure to total molecules vary (Svensson, 2020).

This property allows us to emulate what observed counts in a cell would look like at a lower exposure (total molecule counts per cell) using *binomial thinning*,

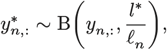

where *l*^*\**^ is as *target depth* for the thinned cell.

### Training SCVI with binomial thinning

To encourage the encoder for the SCVI model to be more robust to low UMI depths we attempt encoding thinned observations while having the decoder aim at reconstructing the observed gene expression counts,

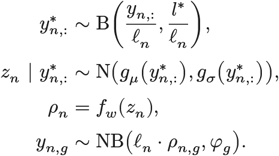

During training, target depths *l*^*\**^ are dynamically sampled for each cell in each minibatch from the log-uniform distribution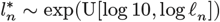.

### Joint embedding-based self-supervised learning

We encode both the original observation and the thinned observation

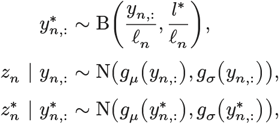

but reconstruct only the original observation,

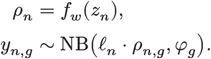

The embeddings of the thinned data is instead used to compute a *cross-correlation loss* within minibatches (Zbontar et al., 2021):

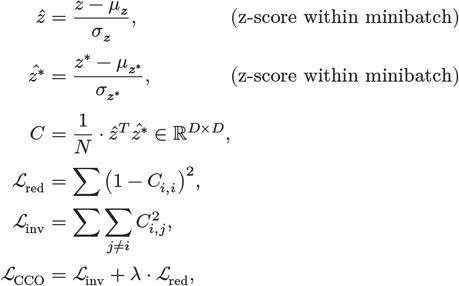

where we use *λ* = 0.01.

The aim of the invariance term ℒ_inv_ is to perfectly correlate the embeddings from the original and thinned versions of the data, while the redundancy term ℒ_red_ encourages decorrelation of different embedding dimensions to avoid collapse to a trivial optimum.

To further prevent collapse we also use a variance loss term,

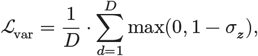

to penalize latent dimensions with low variance (Bardes et al., 2021). The term is averaged across original and thinned views of the cells.

The total loss for this model is

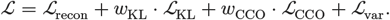

In the ‘joint-embedding only’ ablation we remove the ℒ_recon_ term.

### Measuring representation quality over depth

To investigate the influence of the KL regularization term ℒ_KL_ on representations as a function of UMI depth, we perform binomial thinning on high-UMI cells (*≥* 75th percentile within dataset) and encode the thinned cells. At the thinned target depths, we calculate the distance from the origin of representation space to the representation of the thinned cell, ‖*z*^*\**^‖. The process is performed for a sample of 1,000 high-UMI cells and the analysis is replicated five times before saving the average distance to origin.

We define a learned bias point as the mean representation of a cell with no observed molecule counts,

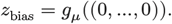

Similar to above, we measure the distance to the bias point *z*_bias_ for representation of thinned data, ‖*z*_bias_ *− z*^*\**^‖. The analysis uses the same sampling strategy as for estimating the average distance to the origin, and is similarly averaged.

In addition to these technical properties, we define metrics for preservation of biological aspects of the data that relate to typical computational biology tasks.

#### Cluster accuracy

Unsupervised clustering is a popular analysis strategy to identify consistent cell state populations. Cluster membership should be preserved for cells with lower UMI depth.

Using representations of high-UMI cells (*≥* 75th percentile within dataset) from the reference SCVI model, unsupervised Leiden clustering defines ‘true’ cluster labels. High-UMI cells with labels are thinned to target depths, then encoded with a given variant of a model to create thinned embeddings *z*^*\**^. Using these thinned embeddings, for each cell, we evaluate if the nearest neighbor of the cell has the same cluster label. The results are quantified by accuracy of nearest neighbor cluster labels. For each experiment we sample 5,000 cells, and the procedure is replicated five times before reporting the average cluster accuracy.

#### Condition accuracy

Differences between cells due to experimental conditions should be reflected in their representations. Preserving differences between experimental at lower depths enable better comparative analysis.

High-UMI cells are labeled by the known experimental or biological condition of the sample they were collected from. After thinned cells are encoded, the representations are used to evaluate the condition label of the nearest neighbor of each cell. For each condition label, 500 labeled cells are sampled and thinned to the given target depths. The average accuracy of condition labels is reported, and the process is replicated five times before saving the average.

#### Sample accuracy

In addition to differences in biological conditions, observed variation between biological samples is often important to investigate, and relate to the magnitudes of differences in biological conditions.

To quantify stability in reflecting sample-to-sample variation, we perform the same analysis as for condition accuracy. However, we use biological sample labels, including biological replicates, instead of condition labels.

### Measuring reconstruction quality

The SCVI model is designed to optimize reconstruction performance under a negative binomial model of observed gene expression counts. Statistical compatibility of counts can be abstract and does not represent all desired properties of good reconstruction. We define several metrics to capture various properties of reconstructions relative to the observed data.

#### Statistical compatibility

Individual studies and datasets measure or analyze different sets of genes, making the ℒ_recon_ unit hard to compare between datasets. To make it easier to compare results between datasets we define the metric ℒ_recon_/*G*, written as ‘NLL per gene’.

#### Separating zero and nonzero observations

In gene expression data the majority of the observations *y*_*n,g*_ are 0 (80-95%). As a consequence, a naive model that predicts very small *µ*_*n,g*_ will have very similar ℒ_recon_/*G* value to a successfully fitted model.

To focus on the more difficult cases where nonzero molecule counts have been observed, we define the measure

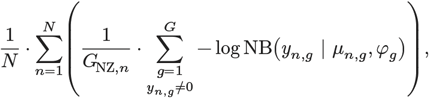

where *G*_NZ,*n*_ is the number of nonzero genes in cell *n* (Palla et al., 2025). We refer to this measure as ‘NLL per gene (NZ)’.

As a complementary measure, we calculate

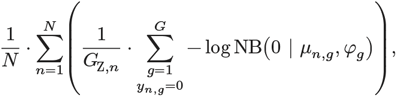

where *G*_Z,*n*_ is the number of zero count observations in cell *n*. We refer to this measure as ‘NLL per gene (Z)’.

#### Interpretable magnitude errors

The negative log likelihoods quantifies the probabilistic compatibility between model predictions and observations, including distribution uncertainty. Unfortunately, it is difficult to relate this unit to magnitudes that are interpreted in practice.

When comparing molecule counts between biological samples, practitioners often look at log2 fold change, log_2_(*x*/*y*). In this prediction setting, for the non-zero portion of the data, we can calculate the same value: log_2_(*µ*_*n,g*_/*y*_*n,g*_). The mean of the absolute values of these log2 fold changes shows us ‘for an average observation, the models predictions are X fold off’,

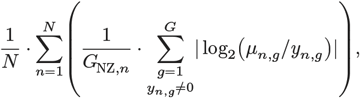

a form of mean average error. We refer to this metric as ‘| log2(μ/x)|’. To detect if model predictions are biased in any direction we calculate the same mean without first taking the absolute value,

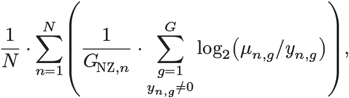

which we refer to as ‘log2(μ/x)’.

#### Gene correlation across cells

Negative log likelihood and log2 fold change measures how good cell, gene -reconstructions are on an absolute scale. An important aspect of model reconstructions is how different gene expression levels are distributed across the cells. This allows distinguishing between cells with high or low expression of a given gene across the dataset.

To quantify whether apparent distributions of observed gene counts are preserved in model reconstructions we use Pearson correlation between observations and predictions across the cells for a given gene.

We calculate Pearson correlation in two different ways. Including all gene counts across the cells, we calculate

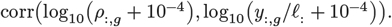

We refer to this metric as ‘r (all)’.

For the non-zero fraction, we calculate

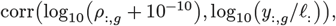

and refer to thos metric as ‘r (nz)’.

## Data

This work focuses on applications of SCVI on typical datasets. A collection of six representative datasets were curated from recent scRNA-seq studies.

### GSE282881

89,321 peripheral blood mononuclear cells from four healthy human donors, measuring expression of 3,086 genes using SCITO-seq2 (Lee et al., 2026). This dataset has no biological conditions to separate, but donor identity was used to assess sample accuracy.

### GSE288481

41,256 Pdgfra+ cells from Il11mNG reporter mouse intestine, measuring 22,313 genes using 10x Genomics Chromium Next GEM Chip G (v3.1). The study compares samples from acute DSS treated and chronic DSS treated mice (Pokatayev et al., 2026). Condition accuracy attempts to separate the DSS treatments, while sample accuracy attempted to separate the five biological mouse samples.

### GSE288761

13,641 nuclei from brains of murine Alzheimer models, measuring expression of 18,730 genes using 10x Genomics Chromium with HTO multiplexing. The study compares samples after conditional activation of a *Braf* V600E allele in *Cx3cr1* expressing cells with controls, in two different brain regions (Vicario et al., 2025). Condition accuracy in this settings classifies by the combinations of induction status and brain region. Sample accuracy classifies the eight biological samples in the dataset.

### GSE308009

39,827 cells from mouse hindlimb knee joints measuring expression of 30,266 genes using Parse Biosciences Evercode WT Mega v1 kit. The study compares the effect of a PGDH inhibitor with a negative control between young and aged mice (Singla et al., 2025). For condition accuracy, the combination of age and treatment status was used. For sample accuracy, all 16 biological samples were used.

### GSE310923

51,277 peripheral blood mononuclear cells from young and old mice measuring expression of 32,338 genes using 10x Genomics Chromium Next GEM Single Cell 3’ Kit (v3.1). The original paper compared the hematopoietic system between young and old mice (White et al., 2026). Condition accuracy classified cells from young or old mice, while sample accuracy classified each of the 12 independent samples.

### OMIX002221

90,852 cells from mouse spleens under induced sepsis compared to negative control and attenuating treatment, measuring 32,285 genes using 10x Genomics Chromium Next GEM single-cell 3′ Kit (v3.1). The study was designed to investigate of artesunate could be used to attenuate sepsis (Chen et al., 2023). Condition accuracy separates negative control, CLP-induced sepsis, and CLP-induced sepsis together with artesunate treatment. Sample accuracy separates the nine independent mice.

## Code

Implementations of the models are available in a branch on github: https://github.com/vals/scVI/tree/scvi-joint-embedding

Scripts for fitting models, and generating figures and tables, are available at https://github.com/vals/scvi-joint-embedding-reproducibility

## Supplementary material

**Figure S1:**
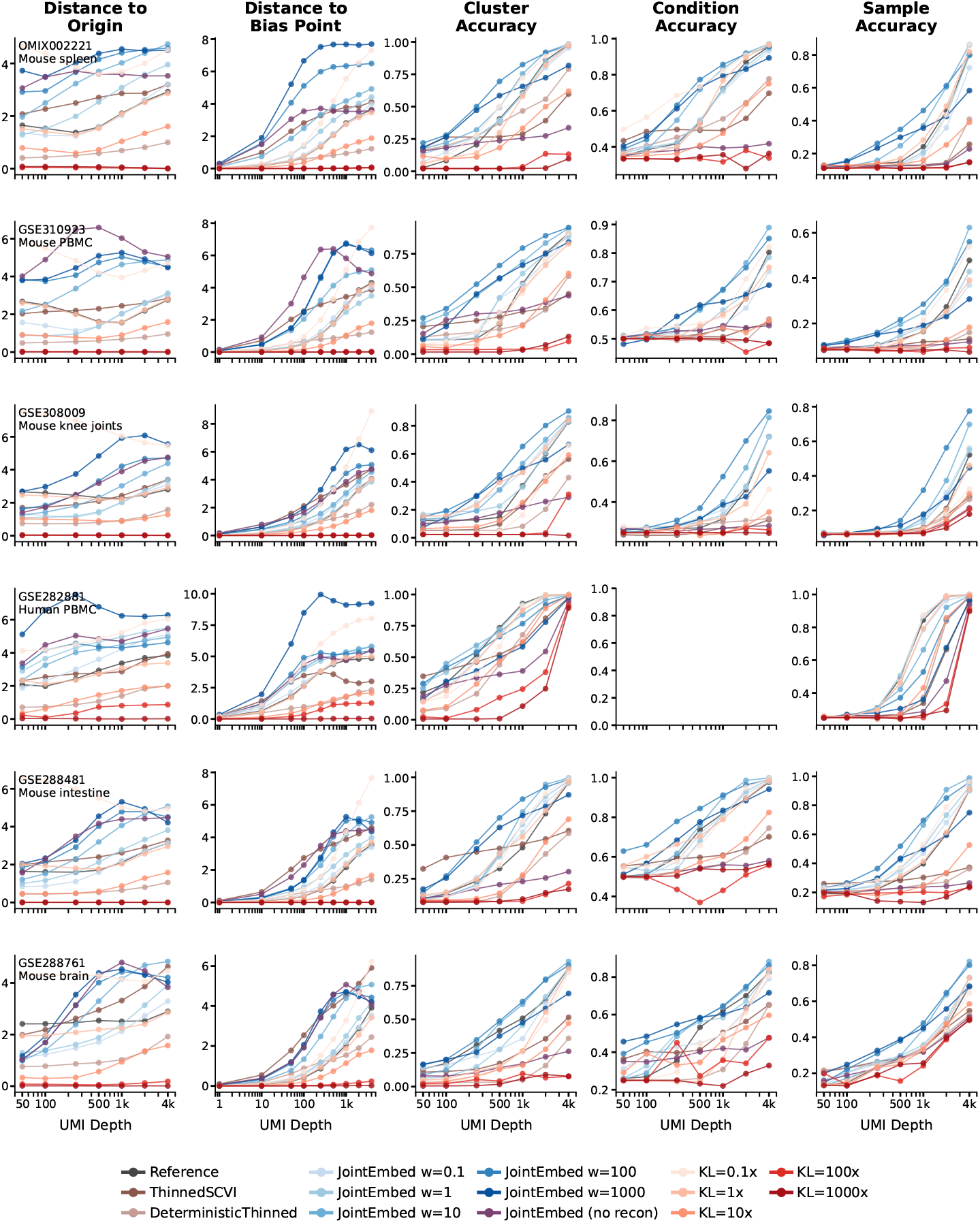
Distance to origin, distance to bias point, and representation performance metrics ‘cluster accuracy’, ‘condition accuracy’, and ‘sample accuracy’ as a function of thinned UMI depth across all models and datasets considered.

**Table S1:** Reconstruction metrics for KL weight ablation variants across all datasets.

| Model | $\vert \log_2(\mu/x)\vert $ | $\log_2(\mu/x)$ | NLL (nz) | NLL (z) | NLL (all) | r (all) | r (nz) |
| --- | --- | --- | --- | --- | --- | --- | --- |
| **OMIX002221** |  |  |  |  |  |  |  |
| Predict 0 | — | — | 48.26 | 0.00 | 2.63 | 0.000 | 0.000 |
| Gene mean × lib | 2.26 | -2.06 | 2.66 | 0.05 | 0.17 | — | — |
| Gene mean (const) | **1.99** | **-1.53** | 2.86 | 0.06 | 0.21 | 0.000 | 0.000 |
| Reference SCVI | 2.12 | -1.96 | 2.38 | **0.04** | 0.15 | 0.092 | 0.209 |
| KL 0.1× | 2.07 | -1.92 | **2.34** | 0.04 | **0.15** | **0.168** | **0.272** |
| KL 1× | 2.09 | -1.93 | 2.36 | 0.04 | 0.15 | 0.152 | 0.261 |
| KL 10× | 2.15 | -1.97 | 2.45 | 0.04 | 0.15 | 0.100 | 0.236 |
| KL 100× | 2.25 | -2.05 | 2.59 | 0.04 | 0.16 | -0.015 | 0.234 |
| KL 1000× | 2.28 | -2.08 | 2.61 | 0.04 | 0.16 | -0.015 | 0.238 |
| **GSE310923** |  |  |  |  |  |  |  |
| Predict 0 | — | — | 48.23 | 0.00 | 4.13 | 0.000 | 0.000 |
| Gene mean × lib | 1.71 | -1.49 | 2.32 | 0.07 | 0.24 | — | — |
| Gene mean (const) | **1.58** | **-1.22** | 2.37 | 0.08 | 0.28 | 0.000 | 0.000 |
| Reference SCVI | 1.60 | -1.39 | **2.15** | 0.06 | 0.22 | 0.105 | **0.206** |
| KL 0.1× | 1.62 | -1.42 | 2.16 | **0.05** | **0.21** | **0.178** | 0.192 |
| KL 1× | 1.60 | -1.40 | **2.15** | 0.06 | 0.22 | 0.161 | 0.203 |
| KL 10× | 1.66 | -1.45 | 2.22 | 0.05 | 0.22 | 0.127 | 0.187 |
| KL 100× | 1.70 | -1.48 | 2.32 | 0.06 | 0.23 | -0.020 | 0.148 |
| KL 1000× | 1.71 | -1.49 | 2.32 | 0.06 | 0.23 | -0.019 | 0.145 |
| **GSE308009** |  |  |  |  |  |  |  |
| Predict 0 | — | — | 61.49 | 0.00 | 5.48 | 0.000 | 0.000 |
| Gene mean × lib | 2.00 | -1.66 | 2.80 | 0.08 | 0.29 | — | — |
| Gene mean (const) | **1.75** | **-1.16** | 2.65 | 0.10 | 0.33 | 0.000 | 0.000 |
| Reference SCVI | 1.88 | -1.55 | 2.61 | **0.06** | 0.26 | 0.148 | 0.257 |
| KL 0.1× | 1.83 | -1.49 | **2.57** | 0.06 | **0.26** | **0.151** | **0.347** |
| KL 1× | 1.87 | -1.54 | 2.61 | 0.06 | 0.26 | 0.136 | 0.326 |
| KL 10× | 1.90 | -1.55 | 2.68 | 0.06 | 0.27 | 0.091 | 0.273 |
| KL 100× | 1.97 | -1.62 | 2.80 | 0.07 | 0.28 | -0.026 | 0.285 |
| KL 1000× | 1.97 | -1.62 | 2.80 | 0.07 | 0.28 | -0.026 | 0.279 |
| **GSE282881** |  |  |  |  |  |  |  |
| Predict 0 | — | — | 46.01 | 0.00 | 6.27 | 0.000 | 0.000 |
| Gene mean × lib | 2.47 | -2.19 | 3.05 | 0.10 | 0.39 | — | — |
| Gene mean (const) | 2.08 | **-1.46** | 3.09 | 0.16 | 0.50 | 0.000 | 0.000 |
| Reference SCVI | 1.81 | -1.53 | 2.30 | 0.07 | 0.30 | 0.273 | 0.359 |
| KL 0.1× | **1.79** | -1.51 | **2.27** | 0.07 | **0.30** | **0.284** | 0.386 |
| KL 1× | 1.83 | -1.57 | 2.32 | **0.07** | 0.30 | 0.267 | **0.393** |
| KL 10× | 1.89 | -1.61 | 2.38 | 0.07 | 0.31 | 0.246 | 0.352 |
| KL 100× | 2.03 | -1.76 | 2.56 | 0.08 | 0.33 | 0.141 | 0.275 |
| KL 1000× | 2.20 | -1.93 | 2.77 | 0.09 | 0.36 | 0.001 | -0.001 |
| **GSE288481** |  |  |  |  |  |  |  |
| Predict 0 | — | — | 37.63 | 0.00 | 5.86 | 0.000 | 0.000 |
| Gene mean × lib | 1.54 | -1.29 | 2.16 | **0.10** | 0.42 | — | — |
| Gene mean (const) | **1.46** | **-1.12** | 2.25 | 0.11 | 0.45 | 0.000 | 0.000 |
| Reference SCVI | 1.50 | -1.26 | 2.09 | 0.10 | 0.40 | 0.135 | 0.148 |
| KL 0.1× | **1.46** | -1.23 | **2.06** | 0.11 | **0.39** | **0.158** | **0.163** |
| KL 1× | 1.48 | -1.24 | 2.08 | 0.11 | 0.40 | 0.143 | 0.143 |
| KL 10× | 1.50 | -1.25 | 2.12 | 0.11 | 0.40 | 0.107 | 0.114 |
| KL 100× | 1.54 | -1.29 | 2.20 | 0.11 | 0.42 | 0.026 | -0.075 |
| KL 1000× | 1.55 | -1.29 | 2.20 | 0.11 | 0.42 | 0.026 | -0.075 |
| **GSE288761** |  |  |  |  |  |  |  |
| Predict 0 | — | — | 29.68 | 0.00 | 4.23 | 0.000 | 0.000 |
| Gene mean × lib | 2.49 | −2.33 | 2.44 | **0.10** | 0.39 | — | — |
| Gene mean (const) | **1.93** | **−1.64** | 2.62 | 0.12 | 0.45 | 0.000 | 0.000 |
| Reference SCVI | 2.13 | −1.96 | 2.44 | 0.10 | 0.36 | 0.167 | **0.297** |
| KL 0.1× | 2.13 | −1.95 | **2.43** | 0.10 | **0.35** | **0.189** | 0.272 |
| KL 1× | 2.12 | −1.95 | **2.43** | 0.10 | 0.36 | 0.165 | 0.281 |
| KL 10× | 2.25 | −2.08 | 2.54 | 0.10 | 0.36 | 0.127 | 0.256 |
| KL 100× | 2.44 | −2.28 | 2.72 | 0.11 | 0.39 | 0.035 | 0.002 |
| KL 1000× | 2.46 | −2.30 | 2.73 | 0.11 | 0.39 | 0.024 | −0.069 |

**Table S2:** Reconstruction metrics for model architecture variants across all datasets.

| Model | $\vert \log_2(\mu/x)\vert $ | $\log_2(\mu/x)$ | NLL (nz) | NLL (z) | NLL (all) | r (all) | r (nz) |
| --- | --- | --- | --- | --- | --- | --- | --- |
| **OMIX002221** |  |  |  |  |  |  |  |
| Predict 0 | — | — | 48.26 | 0.00 | 2.63 | 0.000 | 0.000 |
| Gene mean × lib | 2.26 | -2.06 | 2.66 | 0.05 | 0.17 | — | — |
| Gene mean (const) | 1.99 | -1.53 | 2.86 | 0.06 | 0.21 | 0.000 | 0.000 |
| Reference SCVI | 2.12 | -1.96 | 2.38 | 0.04 | 0.15 | 0.092 | 0.209 |
| Thinned SCVI | **1.85** | **-1.48** | 3.68 | 0.03 | 0.21 | 0.058 | 0.283 |
| Det. Thinned | 1.90 | -1.56 | 3.68 | **0.02** | 0.20 | 0.066 | **0.315** |
| JointEmbed 0.1 | 2.21 | -2.05 | 2.44 | 0.03 | 0.15 | 0.087 | 0.288 |
| JointEmbed 1 | 2.19 | -2.04 | 2.43 | 0.03 | 0.15 | 0.092 | 0.239 |
| JointEmbed 10 | 2.09 | -1.93 | **2.36** | 0.04 | **0.15** | **0.095** | 0.254 |
| JointEmbed 100 | 2.12 | -1.96 | 2.38 | 0.04 | 0.15 | 0.094 | 0.237 |
| JointEmbed 1000 | 2.08 | -1.92 | **2.36** | 0.04 | 0.15 | 0.089 | 0.231 |
| **GSE310923** |  |  |  |  |  |  |  |
| Predict 0 | — | — | 48.23 | 0.00 | 4.13 | 0.000 | 0.000 |
| Gene mean × lib | 1.71 | -1.49 | 2.32 | 0.07 | 0.24 | — | — |
| Gene mean (const) | 1.58 | -1.22 | 2.37 | 0.08 | 0.28 | 0.000 | 0.000 |
| Reference SCVI | 1.60 | -1.39 | **2.15** | 0.06 | **0.22** | 0.105 | 0.206 |
| Thinned SCVI | **1.31** | **-0.86** | 3.67 | 0.04 | 0.32 | 0.075 | 0.236 |
| Det. Thinned | 1.42 | -1.00 | 3.69 | **0.03** | 0.32 | 0.084 | 0.218 |
| JointEmbed 0.1 | 1.75 | -1.57 | 2.25 | 0.05 | 0.22 | 0.105 | 0.253 |
| JointEmbed 1 | 1.75 | -1.56 | 2.25 | 0.05 | 0.22 | 0.106 | 0.196 |
| JointEmbed 10 | 1.66 | -1.47 | 2.19 | 0.05 | **0.22** | **0.107** | **0.272** |
| JointEmbed 100 | 1.62 | -1.42 | 2.17 | 0.05 | 0.22 | 0.104 | 0.224 |
| JointEmbed 1000 | 1.62 | -1.43 | 2.18 | 0.06 | 0.22 | 0.097 | 0.186 |
| **GSE308009** |  |  |  |  |  |  |  |
| Predict 0 | — | — | 61.49 | 0.00 | 5.48 | 0.000 | 0.000 |
| Gene mean × lib | 2.00 | -1.66 | 2.80 | 0.08 | 0.29 | — | — |
| Gene mean (const) | 1.75 | -1.16 | 2.65 | 0.10 | 0.33 | 0.000 | 0.000 |
| Reference SCVI | 1.88 | -1.55 | 2.61 | 0.06 | **0.26** | 0.148 | 0.257 |
| Thinned SCVI | **1.48** | **-0.92** | 4.37 | 0.03 | 0.39 | 0.090 | 0.283 |
| Det. Thinned | 1.66 | -1.08 | 4.40 | **0.03** | 0.38 | 0.103 | 0.314 |
| JointEmbed 0.1 | 1.93 | -1.62 | 2.64 | 0.06 | **0.26** | 0.145 | **0.332** |
| JointEmbed 1 | 1.90 | -1.59 | 2.63 | 0.06 | **0.26** | 0.145 | 0.317 |
| JointEmbed 10 | 1.87 | -1.53 | 2.61 | 0.06 | **0.26** | **0.151** | 0.305 |
| JointEmbed 100 | 1.84 | -1.52 | 2.59 | 0.06 | 0.26 | 0.146 | 0.314 |
| JointEmbed 1000 | 1.80 | -1.45 | **2.58** | 0.07 | 0.27 | 0.127 | 0.227 |
| **GSE282881** |  |  |  |  |  |  |  |
| Predict 0 | — | — | 46.01 | 0.00 | 6.27 | 0.000 | 0.000 |
| Gene mean × lib | 2.47 | -2.19 | 3.05 | 0.10 | 0.39 | — | — |
| Gene mean (const) | 2.08 | -1.46 | 3.09 | 0.16 | 0.50 | 0.000 | 0.000 |
| Reference SCVI | 1.81 | -1.53 | **2.30** | 0.07 | **0.30** | **0.273** | 0.359 |
| Thinned SCVI | **1.70** | **-1.25** | 3.43 | 0.06 | 0.42 | 0.247 | 0.381 |
| Det. Thinned | **1.70** | -1.26 | 3.44 | **0.05** | 0.42 | 0.257 | 0.367 |
| JointEmbed 0.1 | 1.81 | -1.55 | 2.31 | 0.07 | 0.30 | 0.262 | 0.415 |
| JointEmbed 1 | 1.84 | -1.59 | 2.33 | 0.07 | 0.30 | 0.264 | **0.434** |
| JointEmbed 10 | 1.85 | -1.59 | 2.34 | 0.07 | 0.31 | 0.262 | 0.395 |
| JointEmbed 100 | 1.83 | -1.56 | 2.33 | 0.07 | 0.31 | 0.255 | 0.396 |
| JointEmbed 1000 | 1.83 | -1.56 | 2.34 | 0.07 | 0.31 | 0.244 | 0.396 |
| **GSE288481** |  |  |  |  |  |  |  |
| Predict 0 | — | — | 37.63 | 0.00 | 5.86 | 0.000 | 0.000 |
| Gene mean × lib | 1.54 | -1.29 | 2.16 | 0.10 | 0.42 | — | — |
| Gene mean (const) | 1.46 | -1.12 | 2.25 | 0.11 | 0.45 | 0.000 | 0.000 |
| Reference SCVI | 1.50 | -1.26 | 2.09 | 0.10 | 0.40 | **0.135** | 0.148 |
| Thinned SCVI | **1.30** | **-0.95** | 3.61 | 0.06 | 0.59 | 0.077 | **0.328** |
| Det. Thinned | 1.40 | -1.04 | 3.64 | **0.05** | 0.58 | 0.092 | 0.325 |
| JointEmbed 0.1 | 1.53 | -1.30 | 2.11 | 0.10 | 0.40 | 0.131 | 0.194 |
| JointEmbed 1 | 1.52 | -1.29 | 2.11 | 0.10 | 0.40 | 0.131 | 0.192 |
| JointEmbed 10 | 1.49 | -1.26 | 2.09 | 0.10 | **0.40** | **0.135** | 0.212 |
| JointEmbed 100 | 1.46 | -1.23 | **2.07** | 0.11 | 0.40 | 0.130 | 0.199 |
| JointEmbed 1000 | 1.46 | -1.22 | 2.08 | 0.11 | 0.40 | 0.121 | 0.157 |
| **GSE288761** |  |  |  |  |  |  |  |
| Predict 0 | — | — | 29.68 | 0.00 | 4.23 | 0.000 | 0.000 |
| Gene mean × lib | 2.49 | -2.33 | 2.44 | 0.10 | 0.39 | — | — |
| Gene mean (const) | **1.93** | **-1.64** | 2.62 | 0.12 | 0.45 | 0.000 | 0.000 |
| Reference SCVI | 2.13 | -1.96 | 2.44 | 0.10 | 0.36 | 0.167 | 0.297 |
| Thinned SCVI | 2.06 | -1.89 | 3.06 | **0.07** | 0.42 | 0.132 | **0.389** |
| Det. Thinned | 1.94 | -1.80 | 3.00 | 0.09 | 0.43 | 0.148 | 0.197 |
| JointEmbed 0.1 | 2.12 | -1.96 | 2.43 | 0.10 | **0.35** | 0.163 | 0.365 |
| JointEmbed 1 | 2.15 | -1.98 | 2.45 | 0.09 | **0.35** | 0.160 | 0.388 |
| JointEmbed 10 | 2.20 | -2.02 | 2.47 | 0.09 | **0.35** | 0.165 | 0.359 |
| JointEmbed 100 | 2.11 | -1.94 | 2.42 | 0.10 | 0.36 | **0.170** | 0.352 |
| JointEmbed 1000 | 2.03 | -1.87 | **2.39** | 0.11 | 0.36 | 0.158 | 0.311 |

## Footnotes

1 https://www.nxn.se/p/r81gsph58us02le8prfd35nrvfmsvc

2 https://www.nxn.se/p/training-scvi-metal-acceleration

## Bibliography

Aldridge, S., & Teichmann, S. A. (2020). Single cell transcriptomics comes of age. Nat. Commun., 11(1), 4307.

Bardes, A., Ponce, J., & LeCun, Y. (2021). VICReg: Variance-Invariance-Covariance Regularization for Self-Supervised Learning. Arxiv [Cs.cv].

Chen, J., He, X., Bai, Y., Liu, J., Wong, Y. K., Xie, L., Zhang, Q., Luo, P., Gao, P., Gu, L., Guo, Q., Cheng, G., Wang, C., & Wang, J. (2023). Single-cell transcriptome analysis reveals the regulatory effects of artesunate on splenic immune cells in polymicrobial sepsis. J. Pharm. Anal., 13(7), 817–829.

Ding, J., Adiconis, X., Simmons, S. K., Kowalczyk, M. S., Hession, C. C., Marjanovic, N. D., Hughes, T. K., Wadsworth, M. H., Burks, T., Nguyen, L. T., Kwon, J. Y. H., Barak, B., Ge, W., Kedaigle, A. J., Carroll, S., Li, S., Hacohen, N., Rozenblatt-Rosen, O., Shalek, A. K., … Levin, J. Z. (2020). Systematic comparison of single-cell and single-nucleus RNA-sequencing methods. Nat. Biotechnol., 38(6), 737–746.

Gayoso, A., Lopez, R., Xing, G., Boyeau, P., Valiollah Pour Amiri, V., Hong, J., Wu, K., Jayasuriya, M., Mehlman, E., Langevin, M., Liu, Y., Samaran, J., Misrachi, G., Nazaret, A., Clivio, O., Xu, C., Ashuach, T., Gabitto, M., Lotfollahi, M., … Yosef, N. (2022). A Python library for probabilistic analysis of singlecell omics data. Nat. Biotechnol., 40(2), 163–166.

Heumos, L., Schaar, A. C., Lance, C., Litinetskaya, A., Drost, F., Zappia, L., Lücken, M. D., Strobl, D. C., Henao, J., Curion, F., Single-cell Best Practices Consortium, Schiller, H. B., & Theis, F. J. (2023). Best practices for single-cell analysis across modalities. Nat. Rev. Genet., 24(8), 550–572.

Hicks, S. C., Townes, F. W., Teng, M., & Irizarry, R. A. (2018). Missing data and technical variability in single-cell RNA-sequencing experiments. Biostatistics, 19(4), 562–578.

James, L., George, T., Roger, G., & Mohammad, N. (2019). Don’t blame the ELBO! A linear VAE perspective on posterior collapse. Arxiv [Cs.lg].

Kassab, M., Maniero, L., & Beltrame, E. (2026). A hyperparameter benchmark of VAE-based methods for scRNA-seq batch integration. Biorxiv, 705093.

Lause, J., Berens, P., & Kobak, D. (2021). Analytic Pearson residuals for normalization of single-cell RNA-seq UMI data. Biorxiv, 2020.12.01.405886.

Lee, S.-H., Jin, B.-Y., Lee, C.-R., Kim, D. R., Shin, A., Park, S.-G., Kim, Y.-J., Kim, S. H., Choi, M., & Hwang, B. (2026). SCITO-seq2: ultra-high-throughput single-cell transcriptome and epitope sequencing. Genome Biol., 1–20.

Lopez, R., Regier, J., Cole, M. B., Jordan, M. I., & Yosef, N. (2018). Deep generative modeling for single-cell transcriptomics. Nat. Methods, 15(12), 1053–1058.

Luecken, M. D., & Theis, F. J. (2019). Current best practices in single-cell RNA-seq analysis: a tutorial. Mol. Syst. Biol., 15(6), e8746.

P Kingma, D., & Welling, M. (2019). An introduction to variational autoencoders. Found. Trends® Mach. Learn., 12(4), 307–392.

Palla, G., Babu, S., Dibaeinia, P., D Pearce, J., Li, D., A Khan, A., Karaletsos, T., & M Tomczak, J. (2025). Scalable single-cell gene expression generation with latent diffusion models. Arxiv [Stat.ml].

Pokatayev, V., Jaiswal, A., Shih, A. R., Segerstolpe, Å., Li, B., Creasey, E. A., Zhao, Y., Lin, C., Murphy, S., Chou, C.-H., Graham, D. B., & Xavier, R. J. (2026). Bidirectional CRISPR screens decode a GLIS3-dependent fibrotic cell circuit. Nature, 1–10.

Singla, M., Wang, Y. X., Monti, E., Bedi, Y., Agarwal, P., Su, S., Ancel, S., Hermsmeier, M., Devisetti, N., Pandey, A., Bakooshli, M. A., Palla, A. R., Goodman, S., Blau, H. M., & Bhutani, N. (2025). Inhibition of 15-hydroxy prostaglandin dehydrogenase promotes cartilage regeneration. Science, eadx6649, eadx6649.

Svensson, V. (2020). Droplet scRNA-seq is not zeroinflated. Nat. Biotechnol.

Svensson, V., Veiga Beltrame, E. da, & Pachter, L. (2019). Quantifying the tradeoff between sequencing depth and cell number in single-cell RNA-seq. Biorxiv, 762773.

Townes, F. W., Hicks, S. C., Aryee, M. J., & Irizarry, R. A. (2019). Feature selection and dimension reduction for single-cell RNA-Seq based on a multinomial model. Genome Biol., 20(1), 295.

Vicario, R., Fragkogianni, S., Pokrovskii, M., Meyer, C., Lopez-Rodrigo, E., Hu, Y., Ogishi, M., Alberdi, A., Baako, A., Ay, O., Plu, I., Sazdovitch, V., Heritier, S., Cohen-Aubart, F., Shor, N., Miyara, M., Nguyen-Khac, F., Viale, A., Idbaih, A., … Geissmann, F. (2025). Role of clonal inflammatory microglia in histiocytosis-associated neurodegeneration. Neuron, 113(7), 1065–1081.e13.

White, R. R., Xiong, K., Wakai, M., Surian, A., Adler, C., Negron, N., Ni, M., Shavlakadze, T., Bai, Y., & Glass, D. J. (2026). A cellular and transcriptomic atlas of the aged mouse hematopoietic system. Aging Cell, 25(2), e70394.

Zbontar, J., Jing, L., Misra, I., LeCun, Y., & Deny, S. (2021). Barlow Twins: Self-supervised learning via redundancy reduction. Arxiv [Cs.cv].

